# Seafloor video-acoustic monitoring in a Greenlandic glacial fjord records hyperbenthos, backward-swimming fish, and narwhals

**DOI:** 10.64898/2026.02.27.708539

**Authors:** Evgeny A. Podolskiy, Monica Ogawa, Kohei Hasegawa, Makoto Tomiyasu, Shin Sugiyama, Yoko Mitani

**Affiliations:** Arctic Research Center, Hokkaido University, Sapporo, Japan; National Institute of Polar Research, Tachikawa, Japan; Faculty of Fisheries Sciences, Hokkaido University, Hakodate, Japan; Institute of Low Temperature Science, Hokkaido University, Sapporo, Japan; Wildlife Research Center, Kyoto University, Kyoto, Japan

**Author notes:** Corresponding author (EAP).

## Abstract

Autonomous video-acoustic monitoring at the sea-floor can improve our understanding of poorly documented ecosystems and help interpret active or passive acoustic data, but it has been rarely carried out, particularly in the Arctic. This study deployed a video camera synchronized with a hydrophone, combined with red lights and other oceanographic instrumentation, on the bottom of a glacial fjord in Inglefield Bredning, northwest Greenland (to 260 m water depth). Through manual review and automatic analysis of high-frequency images (30 fps) and audios (96 kHz), the conditions and biodiversity near the sea-floor were quantified. The data revealed a highly turbulent environment with abundant suspended particles and fibers, with 88% of 478 detected organisms being Amphipoda, Copepoda, Hydrozoa, and Chaetognatha. Amongst the other observed animals were Decapoda, Liparidae, Pterotracheoidea, Ctenophora, and curious narwhals (*Monodon monoceros*). The number of marine snow particles was highly variable through time and could change up to twofold within several hours. The tide modulated the particle flow direction and speed. Overall, the results show that portable moorings with video recorders are an important tool for exploration of the Arctic seafloor.

## Introduction

Arctic glacial fjords are hotspots of marine life, but they are understudied as a result of their remoteness and difficult access, particularly their seafloor ecosystems. Direct observations near the seafloor can reveal biodiversity and animal behavior and are promising because almost all marine ecosystems studied to date show an increase in biomass in the hyperbenthon (i.e., the benthic boundary layer) relative to the water mass immediately above it [1]. They are also potentially helpful in interpreting other indirect measurements, such as those obtained with active acoustic profilers and hydrophones, which cannot verify the sources of reflections or sounds [2]. However, it is unclear if commonly used monitoring methods are noninvasive and suitable, particularly because both active and passive acoustic devices are known to attract megafauna [3, 4].

As part of a long-term environmental research program since 2012 [5] and an acoustic monitoring program since 2019 [6] at Inglefield Bredning, northwest Greenland, a video camera with red lights and a hydrophone were deployed on the seafloor (260 m water depth) for approximately one week. This experimental approach differs from other commercially available devices that could be used in long-term mooring systems or for hyperbentic sampling [1]. First, it does not deter or attract animals with sound or light, as autonomous platforms with echo-sounders might presumably do [3]. Second and third, in contrast to the high precision Underwater Vision Profiler, UVP [7], or submersible digital holographic cameras, like LISST-Holo2 [8], it records sound and allows monitoring objects larger than macro-zooplankton (i.e., a few centimeters). Finally, it does not sample animals with bottom sledges or traps [1].

Against this background, the objective of this study was to document the seafloor environment, assess its biodiversity, and evaluate the performance of a compact mooring system, including its potential physical interaction with narwhals (*Monodon monoceros*), which has recently been identified as a concern [4]. In particular, we hypothesize that a setup significantly shorter than previously used in the area and equipped only with passive instrumentation (including acoustic and light emitters) might be useful for innovative ecological studies, while being less invasive and less attractive to narwhals and other animals.

## Materials and methods

### Study site, setup, and data

The setup included an underwater camera synchronized to a hydrophone (LoggCAM; Biologging Solutions), a separate sound recorder (SoundTrap ST600, S/N 6229; Ocean Instruments), an acoustic release (Ascent AR; Vemco), a buoy (Viny, 10B-8 with 10.7 kg of buoyancy; Kihoozai-kenkyuujo), an anchor (50 kg of rocks in a net), and a Samson 3/8” line connecting all components into a chain that is ∼2.5 m long (Fig 1). The mooring, excluding the anchor, fits into a Zarges box, weighs less than 15 kg, and is rated to 500 m water depth. The setup was first tested in a pool and shallow waters in Japan [11]. On 1 August 2025 (19:56 UTC), the setup was deployed in Inglefield Bredning, Greenland, approximately between Qeqertaq and Heilprin Glacier, from a boat to a water depth of ∼260 m (drop location: 77°28.120’N, 66°21.411’W). In this paper, all times are given in UTC (local time = UTC−1 h). This location was chosen because it has the highest likelihood of narwhal acoustic presence as compared with other monitoring sites within the fjord [4]. On 9 August 2025 (15:31), the mooring was recovered using a surface deck box VR100-300 with a transponding hydrophone VHTx-69kHz (Vemco) for remote opening of the acoustic release. The work was conducted under a Collaborative Research Agreement with the Greenland Institute of Natural Resources in Nuuk, with access to the field site approved by the Danish Immigration Service.

**Fig 1.**
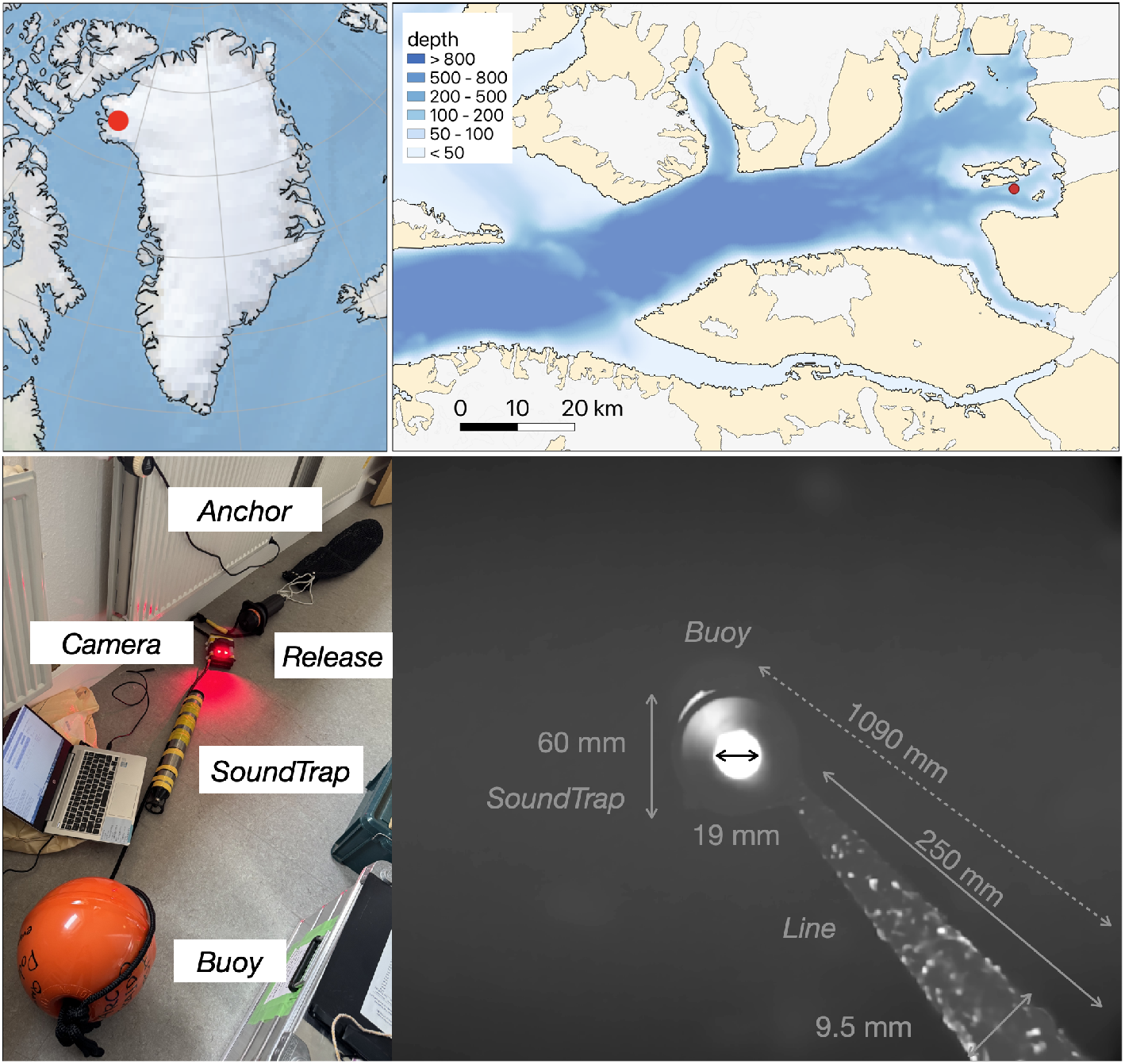
Location of the study site and setup. Inglefield Bredning Fjord, Greenland, with a photograph of the mooring setup and camera view (grayscale, processed, and with any moving particles removed). Lines on the maps delineate study areas and do not necessarily depict accepted national boundaries. The maps were produced using Cartopy and QGIS. Background bed topography and bathymetry are from IceBridge BedMachine Greenland, Version 5 [9, 10]

During the period of deployment, the camera was looking upward and recorded 10-minute-long videos (VGA; 640×480 pixels at 30 frames per second) with audio (96 kHz) every 20 min (i.e., with a 10-min pause between recordings) for about three days. The rationale for the upward viewing direction, as compared with a downward direction that would enable seafloor observations without the accumulation of sediment on the lens, was to film narwhals, which are known to approach from above and hit seafloor moorings [4]. Two LED lights on the camera had a red wavelength (∼660 nm). This choice was made in order to avoid overlap with the retinal absorbance of cetacean eyes [12] and for the presumably passive nature of the observations (i.e., in contrast to standard lights; e.g., [3, 13, 14]). However, due to the high absorbance of the red wavelength, this choice limited the range of observations to <25–100 cm (Fig 1). The SoundTrap was equipped with a pressure relief valve for deep water (Dual Seal PRV Ti - 20 psi; PREVCO Subsea), and continuously recorded sound at a sampling rate of 96 kHz. The internal sensors of the acoustic release recorded the temperature (±0.1°C resolution), pressure (±1 m resolution), average noise, and tilt at 1-min intervals. In summary, seven different streams of data were collected [15–17]. To interpret the water pressure measurements and flow variations, the data were compared with sea-level data from Thule Air Base (renamed Pituffic station), which are collected every minute.

### Data processing

A visual and aural review of all video files was undertaken to detect and identify animals (on a 32-inch Retina 6K Pro Display XDR by Apple), based on previous studies [18–21]. Due to relatively low image quality, caused by red lights and video resolution, taxonomic identification was limited to high taxonomic levels.

Image and audio processing were also undertaken. Existing Matlab functions were used to extract basic video-frame properties. Specifically, for every frame, the total area of the image covered by particles was estimated, the number of particles was counted, and the mean intensity of the color in each image was computed.

Firstly, the background was removed from each processed frame to focus only on moving particles; i.e., by excluding any static, slowly varying features, such as the highly reflective PRV of the SoundTrap bottom, the mooring line, with particles and fibers permanently sticking to it (Fig 1), or sediments on the lens.

Secondly, the background was subtracted from each frame, and the outcome was binarized. This efficiently highlighted the reflective particles and was then used to count the number of particles and integrate their total area. The total area of each frame covered by particles, *A*, is shown as a fraction of the total image size (i.e., *A*/[640 × 480]).

Thirdly, the frame color was treated as follows. True color image data were specified for each pixel of the image as a three-dimensional array of red-green-blue (RGB) triplets. The latter corresponds to a three-element vector, which specifies the intensities of the RGB components of the color as an integer of type “unit8” from [0 0 0] to [255 255 255]. By computing the mean value of each dimension for each frame, the mean intensity of the color was extracted. During the descent of the camera through the water column, each RGB component was computed, and only the temporal variation of red at the seafloor due to the absorption of other colors was considered.

PIVlab was used to study particle motion in the videos. PIVlab is an open-source tool for digital particle image velocimetry (DPIV), which is a common technique in fluid dynamics and is also known as optical flow analysis [22]. To reduce the computational cost, 10 frames were analyzed from each file (0.3 s) instead of processing full videos (150 Gb), and the outcome vectors were averaged for the dominant flow magnitude and direction. Based on the results of the visual review (S1 Table), a MATLAB GUI was used to manually track copepods in the videos (the only animals that repeatedly interacted with the setup).

Finally, both the camera and SoundTrap recorder had hydrophones recording at the same sampling rate. However, because the camera had 10-minute pauses and recorded only for three days, while the SoundTrap was sampling continuously and for the full duration of the experiment, only the SoundTrap acoustic data were visualized, using a long-term spectrogram.

## Results

### Overview

The key variables measured during the camera deployment are shown in Fig 2. The water depth oscillated around 260 m with the tide (amplitudes up to 1.5 m). Measurements started at the minimum tidal range (neap tide), and ended before the maximum tidal range (spring tide); consequently, the camera observations corresponded to the weakest tidal conditions. The mean temperature of the water was -0.18°C. The tilt of the release was constant (3.6°), consistent with the videos that showed no motion or vibration of the setup. The mean background noise level was 169 mV, with spikes corresponding to narwhal ultrasonic vocalizations that are visible on the spectrogram of the SoundTrap data. Narwhals were acoustically present every day, except 6 August 2025 (observed as elevated ultrasound energy at >20 kHz; Fig 2). At lower frequencies, the primary geophonic source was the rumbling, cracking, and melting of icebergs (<10 kHz). Boat engine noise was also recorded (<5 Hz).

**Fig 2.**
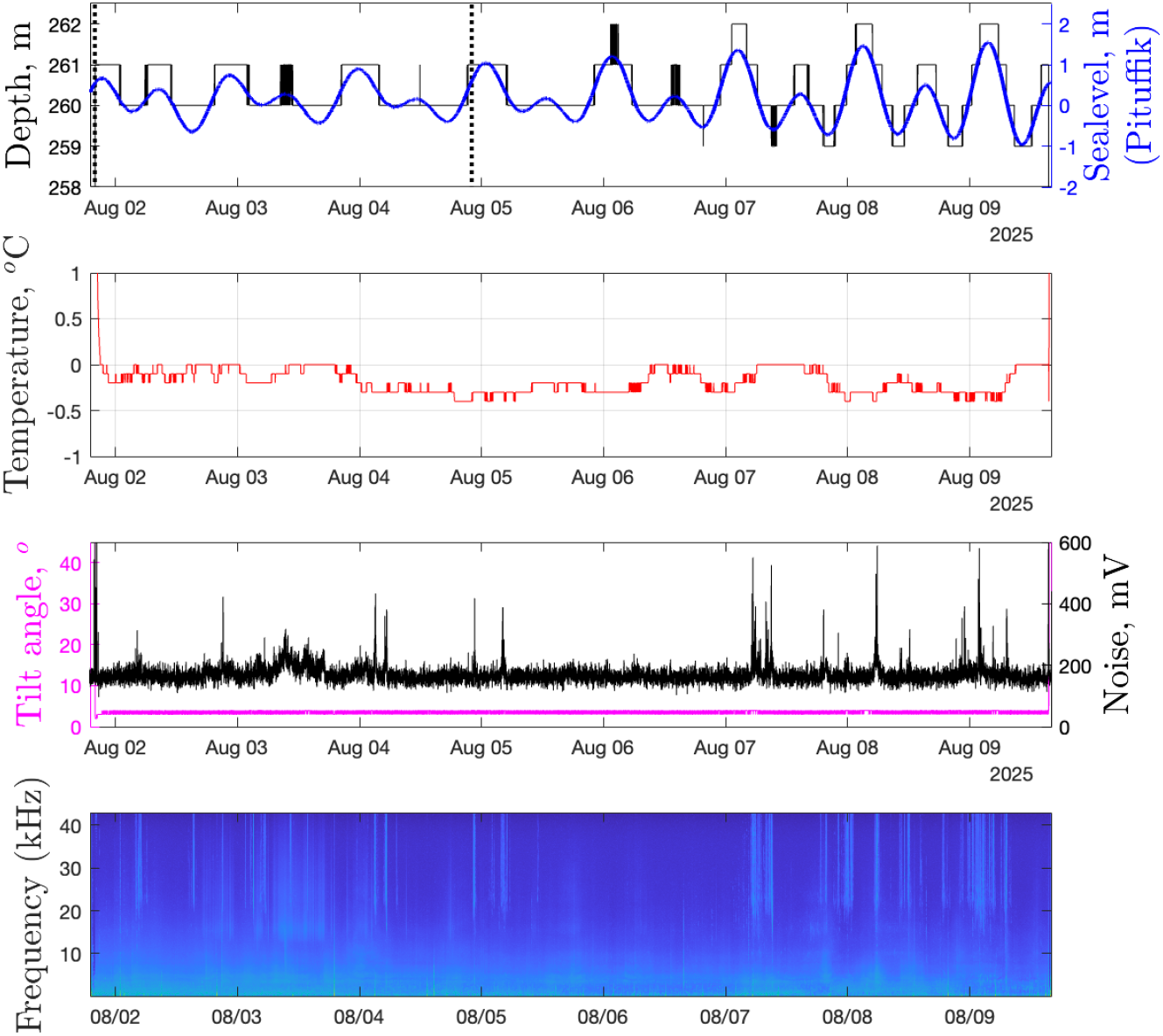
Measurements. Variables measured by the internal sensors of the release and SoundTrap recorder (lower panel) as compared with sea-level data from Thule/Pituffic. Vertical dotted lines mark the interval of the camera operation (i.e., video and sound). The long-term mean spectrogram was computed between 20 and 43,000 Hz, with a time resolution of 10 s and 1024 FFT size.

### Camera

#### Animals

By manually reviewing all 223 video files (37 h in total), we identified actively swimming Amphipoda, heteropods (Pterotracheoidea), arrowworms (Chaetognatha) and bristle worms (Polychaeta), abruptly jumping Copepoda, jellyfish (Hydrozoa), comb jelly (Ctenophora), fish, shrimp (swimming Decapoda), Mysida, and other not identified animals (Fig 3; S1 Table; S1 Video). The organisms were never observed in groups, and usually floated individually, except for some amphipod pairs. In addition, none of these organisms was associated with simultaneously recorded sounds, except for a shrimp scratching the camera and a narwhal. Narwhals were the main source of biophonic sounds, although they did not interact with the mooring and appeared in camera view only once, on 5 August (05:23:13), when high-amplitude narwhal sounds co-occurred with the slow motion of a tusk tip in the background, which was tens of centimeters from the buoy (S1 Video).

**Fig 3.**
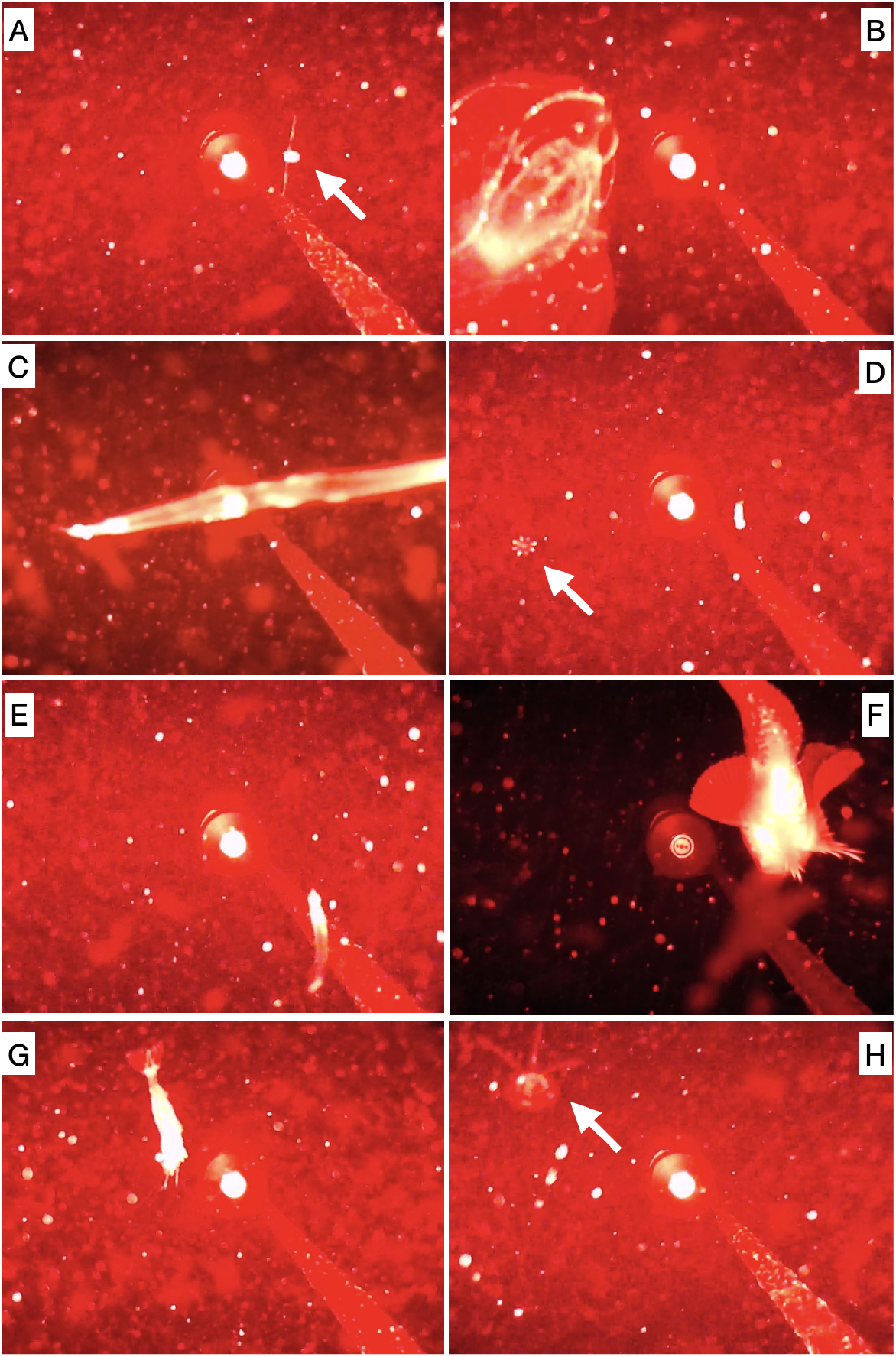
Examples of observed organisms in front of the camera, with the mooring line and SoundTrap in the background. (A) copepod, (B) comb jelly (Ctenophora), (C) arrowworm (Chaetognatha), (D) unidentified organism, (E) juvenile fish, (F) snailfish (Liparidae), (G) shrimp (Decapoda), and (H) jellyfish (Hydrozoa).

The total number of detected specimens was 478 (differentiated in at least 11 taxa), corresponding to approximately 1 detection every 5 minutes. The detection frequency for each category of animal is shown in Fig 4. Amphipoda had the highest occurrence (47%), followed by Copepoda (26%), Hydrozoa (8%), and Chaetognatha (8%; with only one bristle worm, Polychaeta). The remaining ∼12% of detections were mainly unidentified organisms and also fish, Decapoda, heteropods, and Ctenophora. The detection times for the most abundant organisms did not show any clear pattern (Fig 4).

**Fig 4.**
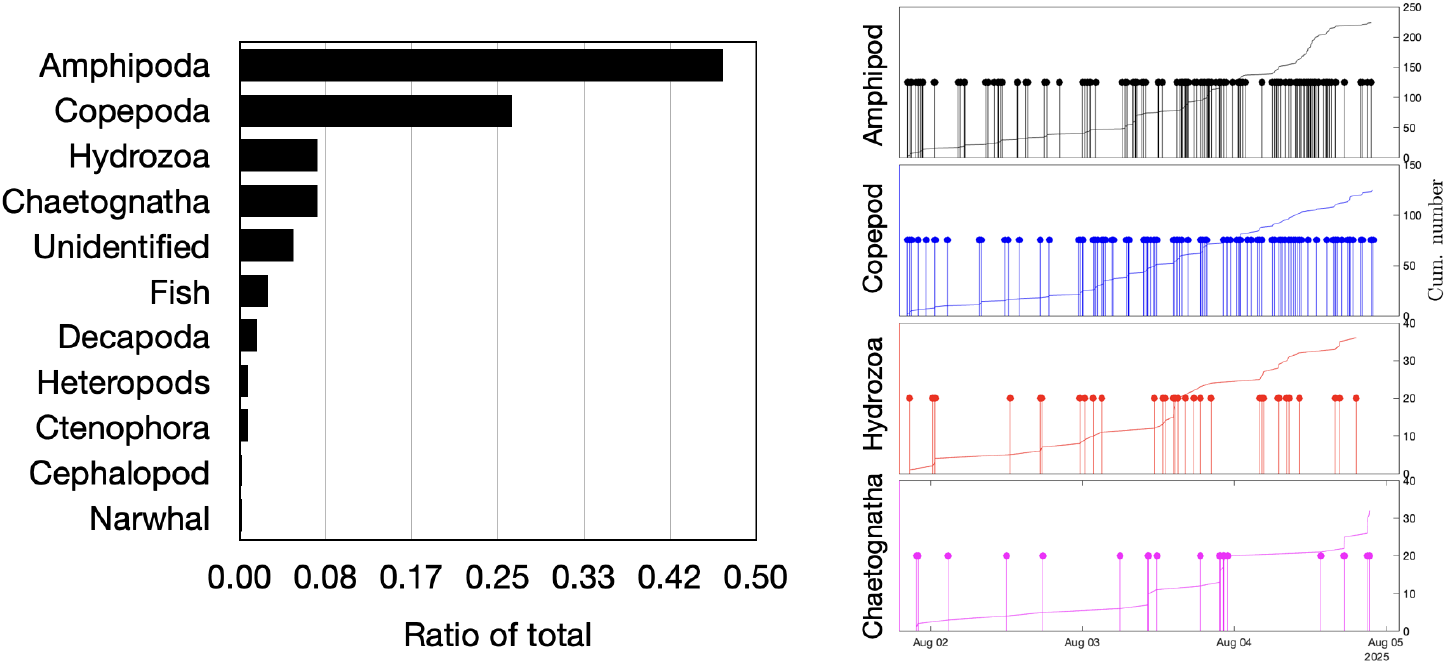
Animal visual detections. Relative numbers of recognized animals and the time of detection of the most frequently observed animals.

The copepods are presumed to be *Calanoida* due to their relatively large size. Specifically, in the frames of some videos (e.g., Nos. 2, 95, and 106), Copepoda collided with the mooring line, for which we know the diameter, indicating antennulae (the first antennae, A1) lengths of up to a centimeter. Interestingly, at the time of collision, the copepods jumped away from the mooring line while drawing their antennulae along their body sides. Such jumpy propulsion was also observed without any obvious contact with the setup (Fig 5).

**Fig 5.**
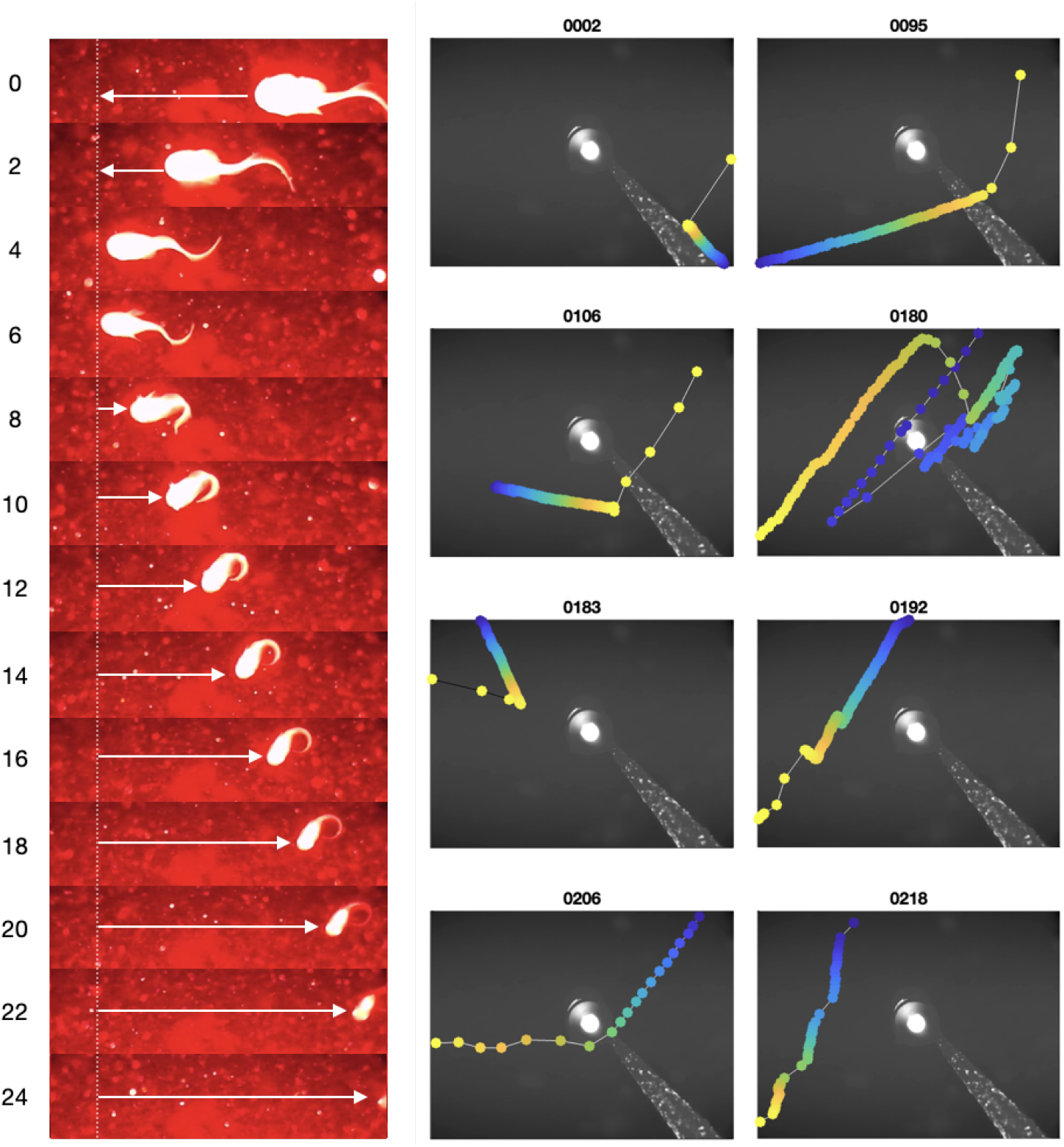
Examples of notable animal motion. (left) Time-lapse sequence of backward swimming by a snailfish at 260 m beneath the surface of the Inglefield Bredning Fjord, Arctic Ocean. The arrows show the direction of movement. Lapsed seconds are shown on the left. Tail folding occurred around the seventh second. Video 142 (frames 13620:60:14340). (right) Frame-by-frame tracking of a copepod jumping movement. The title shows the number of the corresponding video file; the color changes from blue to yellow with time.

The highly active arrowworms, which show characteristic undulatory locomotion [23], were most likely *Eukrohnia hamata*. Clearly filmed fish were a *Liparidae* (also called snailfish in the literature [24]). One snailfish showed peculiar backward swimming, passively drifting backward with the current (Fig 5). It curled its tail and remained motionless for at least 16 s before disappearing from view. The shrimps were likely *Decapoda Thoridae*. Both Liparidae and Decapoda Thoridae are known to be narwhal prey in the fjord [25].

#### Mooring drop and light penetration

After being dropped from the boat, the setup reached the seafloor in 2 min 21 s, which corresponds to a mean rate of descent of 1.84 ms^−1^ . Assuming a constant rate of descent, the change in light conditions can be mapped versus depth with a vertical resolution of ∼6 cm. To achieve this, the mean color intensity was tracked over the course of the drop by computing the mean RGB levels for each frame (Fig 6). This revealed that conditions deeper than 80 m become similar to those on the seafloor, characterized by a minimum amount of green and blue, with numerous suspended particles reflecting the red LED illumination.

**Fig 6.**
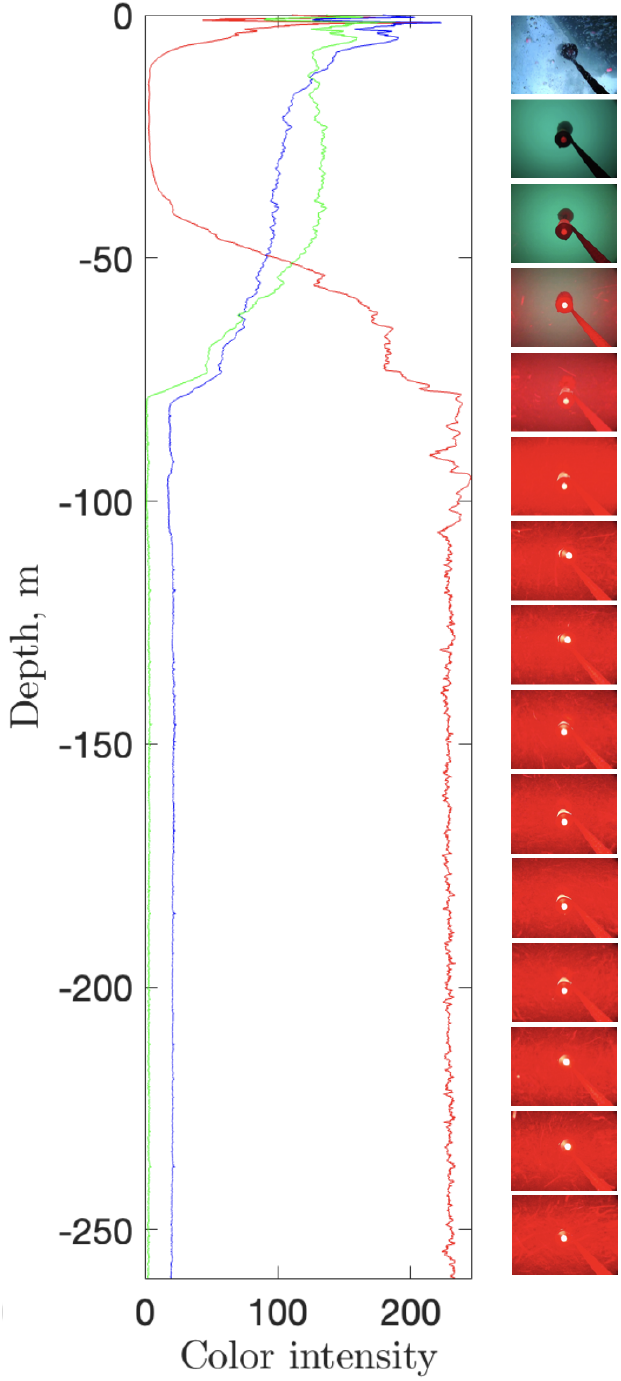
Color variation at deployment of the setup. Variation of mean color intensity during deployment of the setup, translated into depth below sea-level.

#### Suspended matter

At the seafloor, the video revealed numerous small particles and hair-like fibers of varying size, which we are referred to as “marine snow” and assumed to be of organic or mixed origin. Dense fiber clouds were also observed, spinning in front of the camera. These were sometimes caused by organisms, such as fish or shrimp passing nearby (S2 Video). In general, marine snow was constantly moving across the scene. Within a single video file (10 min long), the direction, number, and speed of motion of particles varied substantially. Usually, the motion was translational (i.e., with particles moving horizontally past the camera). However, occasionally, vertical and vortical motion was also evident when particles precipitated, advected away from the lens, or showed apparent shear, with closer and farther particles moving in opposite directions.

Automatically extracted features are presented in Figs 7 – 8. The proportion of each image covered by particles and the absolute number of particles varied substantially with time (Fig 7A,B). The mean proportion of each image covered by particles was 0.95±0.8% (STD), while the mean number of particles was 85±22 (STD). The intensity of red light in each image (Fig 7C) was controlled by the reflecting particles, with three types of transient anomalies resulting from the brightness spike of the first frame (i.e., taken immediately after turning the camera on), animals attaching to the lens, and sediment that had accumulated temporarily on the lens (Fig 7D-E). Power spectral density analysis did not reveal clear cyclicity in the temporal variation of the extracted features (S1 Figure).

**Fig 7.**
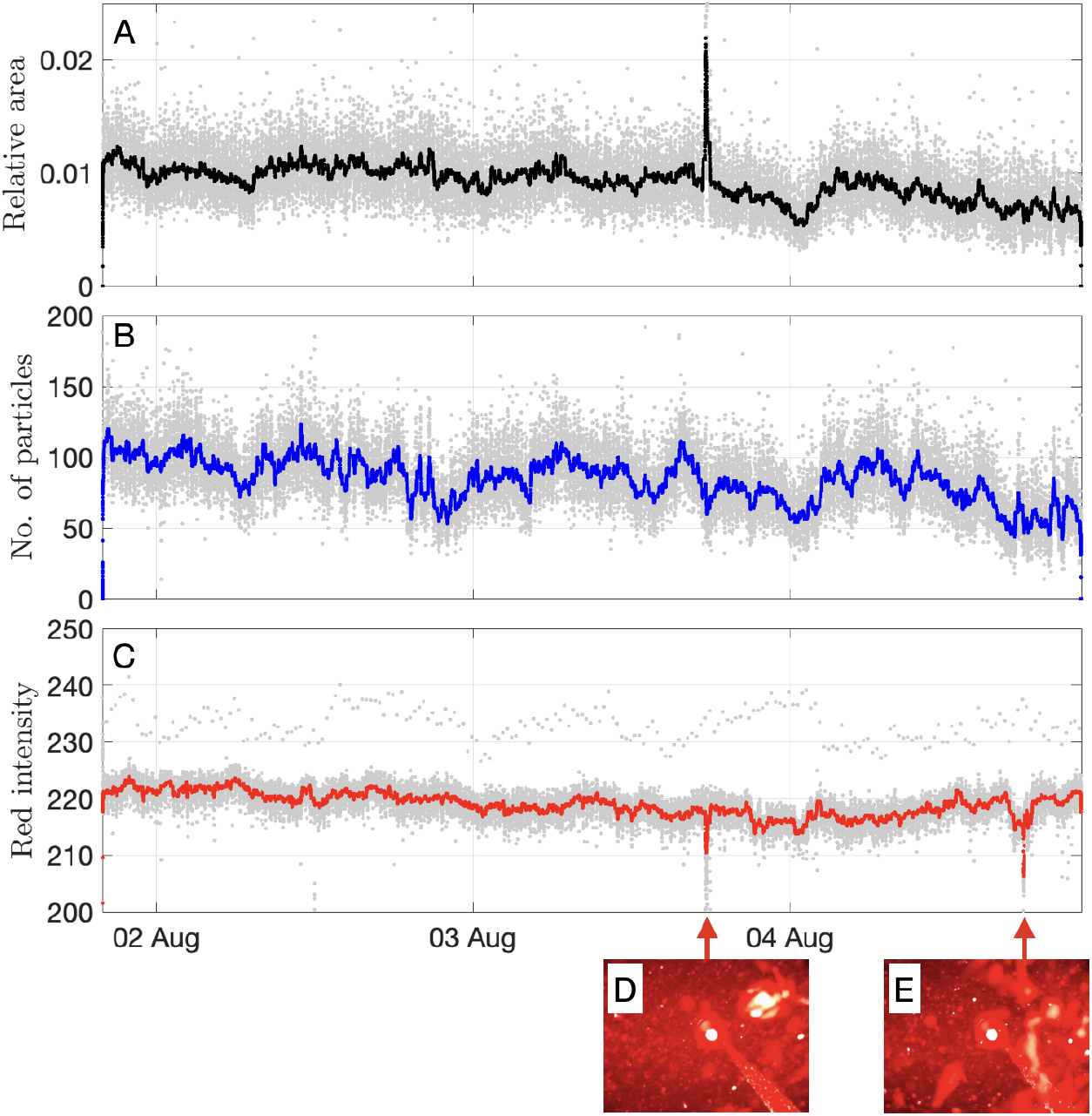
Automatically extracted image features. (A) Relative area of each image covered by detected particles. (B) Number of particles. (C) Mean intensity of the red color. Bold curves correspond to median-filtered time-series using a 3-h sliding window. (D-E) Transient anomalies caused by an organism and sedimentation glow on the lens.

**Fig 8.**
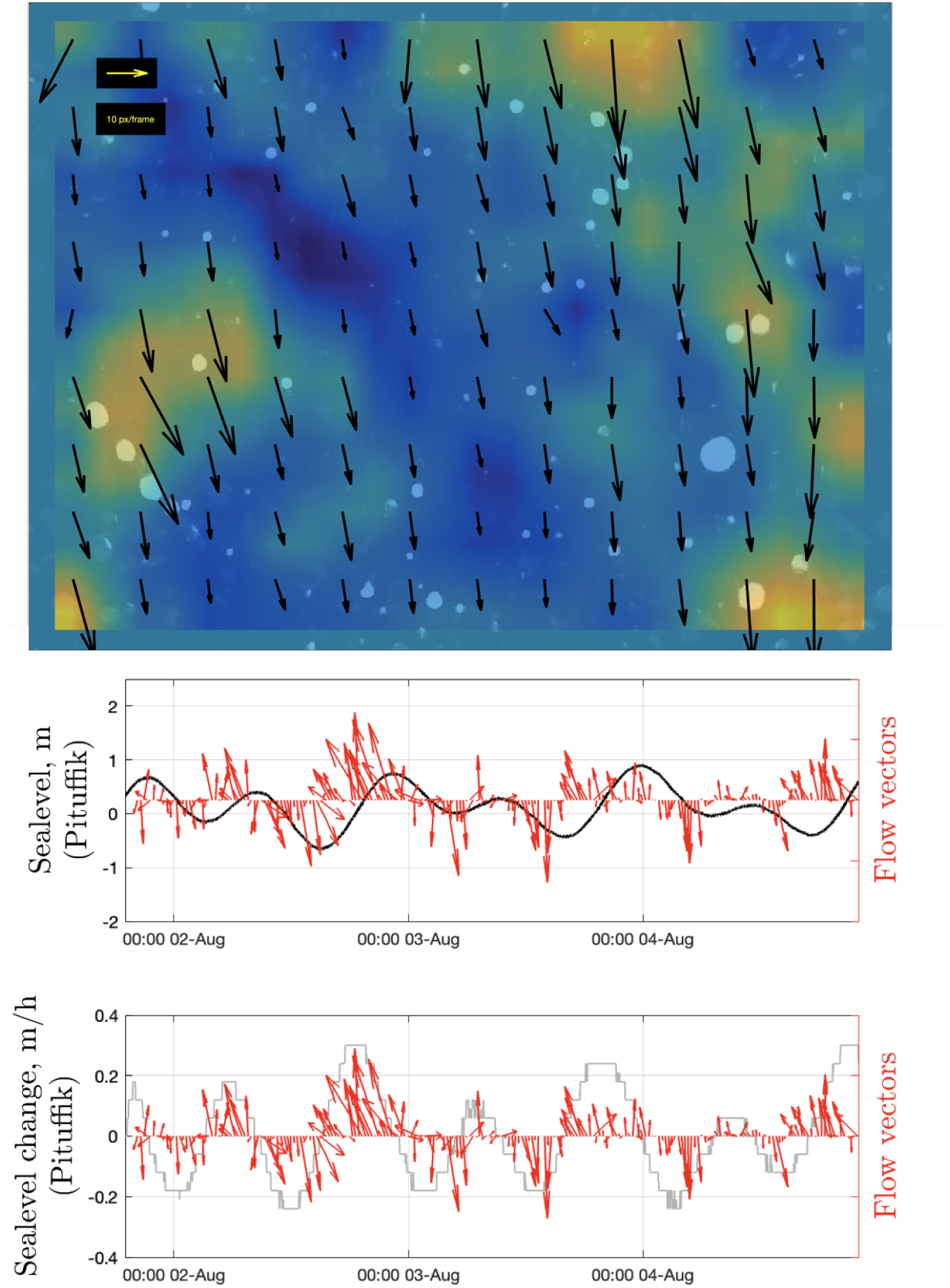
Particle motion. (upper panel) Example of a DPIV result showing the rate and direction of particle flow (the color shows the spatial change in the flow speed). Note that only moving particles can be observed after static background removal. (middle and lower panels) Temporal variations in the particle flow vectors versus sea level and the rate of change of sea level (smoothed with a median filter using a 2-h-long sliding time window).

While the movement direction and speed of individual particles between two sequential frames could be highly variable, the averaged data showed clear 12 h oscillations in these variables (Fig 8; S3 Video). A comparison with the tidal data suggests that the direction of particle motion reverses at high and low water (presumably around slack water). The Pearson’s linear correlation coefficient, *r*, between the sea-level change and the angle of particle motion was 0.54 (*p*-value= 4 × 10^−14^). Moreover, particles moved at the highest speeds during the periods with faster change in water level (Fig 8; r = 0.57; *p*-value= 2 × 10^−16^). This highlights the role of tidal currents in particle transport near the seafloor. Given that the rising tide brings water into the fjord, this implies that the upper edge of each image faces the inner parts of the fjord and the lower edge faces oceanward.

## Discussion

During the three days of camera observations, there were no physical interactions between large objects (such as narwhals) and the mooring, despite their daily acoustic presence. It is unclear whether this reflected the short duration of the experiment or the desired lack of curiosity about the mooring system that was ten times shorter than previously used [4]. The extent of narwhal sensitivity to red light from the camera (660 nm) remains uncertain, given the difficulty of conducting such experiments and the lack of experimental validation. However, it is unlikely that the narwhals were deterred by the camera lights, because retinal pigments of other deep-diving odontocetes are not sensitive to such wavelength [12,26], and their sensitivity is shifted towards blue light, similarly to mesopelagic-dwelling fish [27].

No aggregations of fish or other animals were observed around the mooring. As such, the mooring was not a refuge, and thus was unlikely to have been a foraging attractor for narwhals [4]. The apparently weak aggregation effect is interesting, because fish-aggregating devices, FADs, are commonly used to feed fish and shelter (https://www.fisheries.noaa.gov/national/bycatch/fishing-gear-fish-aggregating-devices). The FADs are floating devices at the surface, and it is possible that floating objects are less effective at close range to the bottom. We acknowledge that the short deployment period may have led to this outcome.

Most of the observed animals could be identified thanks to previous regionally relevant publications [18–21], and stomach content analysis by the authors [25,28]. The explored community corresponded to the hyperbenthos, a term referring to the association of small animals inhabiting the water layer adjacent to the seafloor [1]. As there is no literature on the hyperbenthos in the study area, and low taxonomic resolution is not conducive to detailed comparison with other regions, we limit our brief discussion to general points, functional relationships, and sampling methodology. Amphipods, copepods, medusae, chaetognaths, and decapods are typical residents of the hyperbenthos. Mysida, however, considered a major component of this community and used to detect degradation in response to chemical contaminants [29], was observed only once. Many fish and crustaceans prey on the hyperbenthos and have a hyperbentic life style at early life stages [1]. Commonly, these communities are sampled using bottom sledges and nets, implying that our study provides an alternative methodological approach that may be less reliable for taxonomic identification but yields insights into highly mobile animals in their natural environment.

The high detection rate of Amphipoda and their relatively rapid swimming behaviour near the seafloor is consistent with their scavenging role underwater [4]. The escape reaction of copepods (Fig 5) has previously been studied in the laboratory using high-speed cameras [30], but such direct visual observations of free-swimming copepods in deep waters are rare. The escape reaction appears to primarily represent predator-avoidance behavior triggered by the mechanical sensitivity of the first antennae. Light is an unlikely trigger, as many copepods drifted through the light cone without any response.

Liparidae are vertebrates best adapted to depth and the deepest known fish (down to 8,145 m; [13]). In contrast to the backward swimming by reverse undulation observed for other deep-sea fishes [14], one Liparidae drifted with the current in the present study (Fig 5). Similar behavior has been reported for Liparidae off the coast of central California [24]. Given that this fish does not have a gas bladder, it is not expected to be detectable by active acoustic devices [2].

Subglacial meltwater discharge forms near-surface plumes in front of glaciers that are foraging hot-spots for birds and seals, due to the entrainment of deep water towards the surface [31]. In the present study, the deployment was not made exactly at the calving front of the glacier; however, the deployment depth and negative water temperatures are similar to those at the foot of the previously studied Bowdoin Glacier, which is also located in Inglefield Bredning Fjord [31, 32]. Therefore, considering the exceptionally strong upwelling (which is continuous in summer) and a high likelihood of an osmotic shock [31], it is reasonable to suggest that the species identified in the videos might be carried upward by subglacial plumes and become easy prey. For example, stomach content analysis has revealed that Amphipoda, Liparidae, and Decapoda are preyed upon by seals in the area, while Copepoda are likely prey for sea-birds and ringed seals [28]. Copepoda are the only category that can enter the hyperbenthos from above and return to the water column daily. Due to insufficiently precise taxonomic resolution, it remains unclear whether copepods, amphipods, and shrimps in the hyperbenthos represent different species from those in the stomachs and the water column. However, during the open-water period, four ringed seals tagged with satellite-relay data loggers spent most of their time in the vicinity of tidewater glacier fronts at depths shallower than 100 m [33], further suggesting that they might benefit from foraging opportunities near the surface.

The DPIV revealed tide-modulated changes in flow speed and direction over time, as well as turbulent vortices, highlighting the highly active seafloor conditions (Fig 8). Moreover, a detailed video review indicated that some particle vortices (and thus water mixing) were produced by passing animals (S2 Video, S3 Video), which could further obscure or mask the dynamics at larger scales. There was no clear diurnal or semidiurnal trend in the other extracted parameters (e.g., the number of particles and animals; Figs 4, 6, and S1 Figure), presumably due to the small volume of monitored water, short duration of the video data (3 days), and weak tidal phase during the observation period. This short duration and a lack of observations during spring tide conditions temporally limit conclusions about daily vertical migrations and tidal effects. Nevertheless, given that the clear tidal modulation of particle motion was detected during neap tide conditions, it is reasonable to suggest that spring tide observations might yield a stronger modulation of the considered parameters.

For areas with a high rate of sedimentation, such as near glaciers or rivers [32], the upward orientation of the camera might be disadvantageous and lead to a permanently obscured lens. In such circumstances, a downward-looking setup might be more suitable, although the images might be saturated in extremely turbid waters.

A manual review of video data is a very time-consuming. To detect an organism, their swim patterns were as important as their shape. Going back and forth between the frames while discussing the target with up to three co-authors experienced in species identification required approximately twice as much time as the video duration itself, and corresponded to at least 74 working hours. In this study, a manual review was needed to label and understanding the data. However, in future, a machine vision workflow should be explored for processing large data sets. The labeled dataset might be useful as training data for such a purpose (S1 Table).

Due to the high red-light absorbance and low image resolution, the camera had a limited field of view. On the one hand, our reliance on red light could correspond to a lower likelihood of interference with animal behavior (e.g., Copepoda react to normal light [30]). On the other hand, this might reduce the detection probability and introduce bias. Moreover, poor visibility might result in passively flowing creatures being overlooked or misidentified as particles. For example, in some videos (0141*.mov; >05:42), the amphipod ceased movement and continued to flow with the current. Furthermore, the numerous fibers might be jellyfish tentacles. This suggests that some organisms could be missed, especially during high-speed flow. Therefore, it is appropriate to treat the reported organism counts as detections rather than as abundance estimates. This might be remedied by a higher video resolution.

## Conclusion

Overall, the examples presented above highlight the potential of our approach for innovative environmental and biological studies at the seafloor. So far, there have been few direct underwater observations in the Arctic for ecological research. With video setups becoming accessible [34], more studies would be beneficial for filling this knowledge gap. At this stage, the short-term setup of the present study, which fits into one Zarges box, is suitable for rapid deployments and comparative studies in various locations.

## Supporting information

**S1 Video. Video with examples of passing organisms and their sounds if any**.

**S2 Video. Example of frame-by-frame image processing for computing the number of particles, the corresponding area, and average light intensity**.

**S3 Video. Example of DPIV analysis (for video 0049*.MOV).**

**S1 Table. An annotated table of detected organisms**.

**S1 Figure. Periodograms with 95%-confidence bounds for extracted image features**. The black curve corresponds to the relative area of each image covered by detected particles; the blue curve corresponds to the number of particles, and the red curve corresponds to the mean intensity of the red color.

## Acknowledgments

We thank Y. Sakuragi, I. Qaerngaaq, M. Kristiansen, T. Oshima, and K. Petersen for their assistance with fieldwork logistics. We also thank T. Hirata, D. Lindsay, and J.B. Thiebot for discussions about the underwater filming and light, and T. Koizumi and T. Noda for preparing and customizing the camera. The Editor, Dr. Anish Kumar Warrier, Dr. Riwan Leroux, Dr. Genuario Belmonte, and the anonymous reviewer helped to improve the initial manuscript.

