## Supplementary material for "Seafloor video-acoustic monitoring in a Greenlandic glacial fjord records hyperbenthos, backward-swimming fish, and narwhals": Figure S1

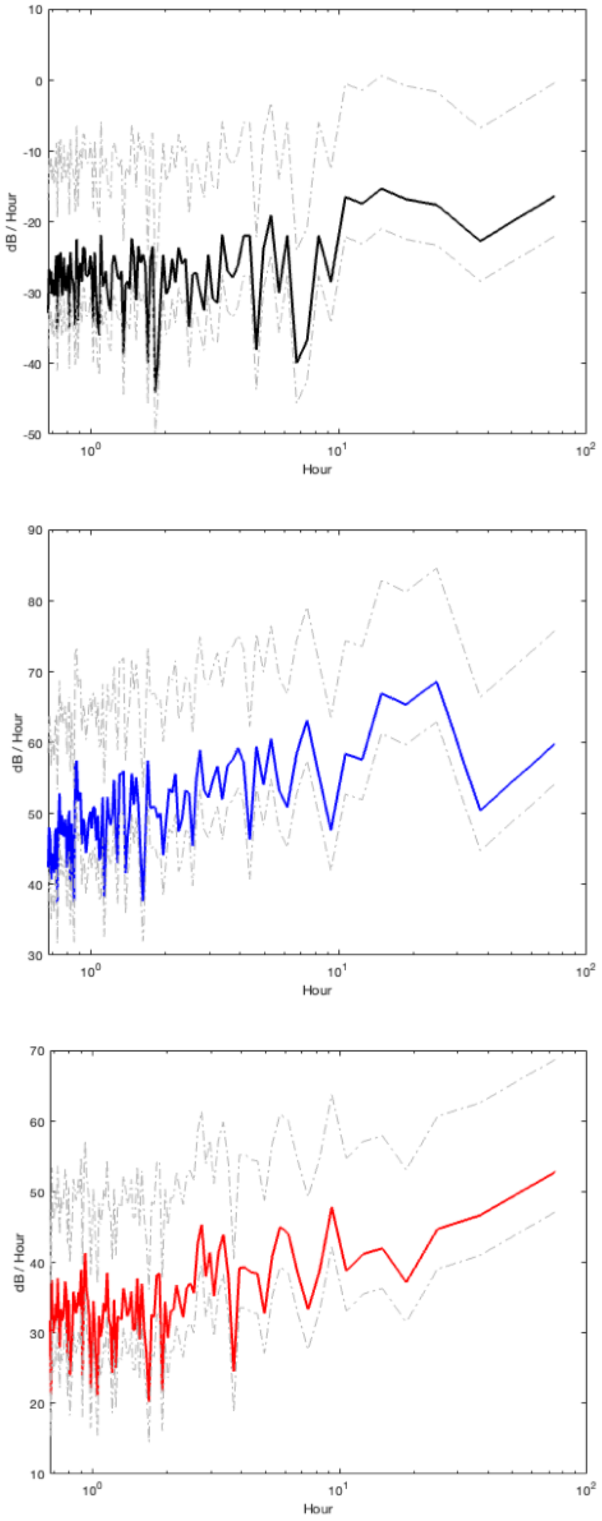


**Fig S1. Periodograms with 95%-confidence bounds for extracted image features.** The black curve corresponds to the relative area of each image covered by detected particles; the blue curve corresponds to the number of particles, and the red curve corresponds to the mean intensity of the red color.
